# Identification of a native novel oncolytic immunoglobulin on exfoliated colon epithelial cells: A bispecific heterodimeric chimera of IgA/IgG

**DOI:** 10.1101/140673

**Authors:** George P. Albaugh, Sudhir K. Dutta, Vasantha Iyengar, Samina Shami, Althaf Lohani, Eduardo Sainz, George Kessie, Prasanna Nair, Sara Lagerholm, Alka Kamra, J-H Joshua Chen, Shilpa Kalavapudi, Robert Shores, Laila E. Phillips, Ram Nair, Padmanabhan P Nair

**Affiliations:** From the Beltsville Human Nutrition Research Centre, Agricultural Research Service, Beltsville, MD 20705; Department of International Health, Johns Hopkins University Bloomberg School of Public Health, Baltimore, MD 21205; Department of Pediatrics, University of Maryland School of Medicine Baltimore, MD 21201; Division of Gastroenterology, Sinai Hospital of Baltimore, Baltimore MD 21215; NonInvasive Technologies LLC, 8170 Lark Brown Road, Suite 101, Elkridge, MD 21075 USA

**Keywords:** Colon epithelial cells, CXCR-4, IgA/IgG chimeric immunoglobulin heterodimer, COX-2, LGR-5, Musashi-1, dedifferentiation, cellular engraftment, oncolysis, gastrointestinal progenitor stem cells (GIP-C)

## Abstract

Understanding the nature of cell surface markers on exfoliated colonic cells is a crucial step in establishing criteria for a normally functioning mucosa. We have found that colonic cells isolated from stool samples (SCSR-010 Fecal Cell Isolation Kit, NonInvasive Technologies, Elkridge, MD), preserved at room temperature for up to one week, with viability of >85% and low levels of apoptosis (8% - 10%) exhibit two distinct cell size subpopulations, in the 2.5μM– 5.0 μM and 5.0μM-8.0μM range. In addition to IgA, about 60% of the cells expressed a novel heterodimeric IgA/IgG immunoglobulin that conferred a broad-spectrum cell mediated cytotoxicity against tumor cells. In a cohort of 58 subjects the exclusive absence of this immunoglobulin in two African-Americans was suggestive of a germline deletion. Serial cultures in stem cell medium retained the expression of this heterodimer. Since a majority of the cystic cells expressed the stem cell markers Lgr5 and Musashi-1 we termed these cells as gastrointestinal progenitor stem cells (GIP-C**). CXCR-4, the cytokine co-receptor for HIV was markedly expressed. These cells also expressed CD20, IgA, IgG, CD45, and COX-2. We assume that they originated from mature columnar epithelium by dedifferentiation. Our observations indicate that we have a robust noninvasive method to study mucosal pathophysiology and a direct method to create a database for applications in regenerative medicine.

---

The human colon renews its mucosa almost every five days and allows the colonization of the lumen by a host of symbionts, microorganisms referred to collectively as the colonic microbiome. This rapid and constant renewal of the mucosa requires a steady source of proliferating cells to maintain a stable turnover rate. This knowledge is attributed to the pioneering work of Leblond and associates, (1) and was later expanded upon by Lipkin and others (2-4).

During early development, cells lining the colonic mucosa acquire the ability to tolerate foreign antigens generated by commensal microflora along with a mixture of antigens of dietary origin. This is made possible through a process of immunological acquiescence resulting in a dynamic equilibrium, which requires the cells of the mucosa to exhibit an adaptive tolerance to the microflora unique to each individual. This process is central to the symbiotic relationship between the host organism and its microflora, (5,6) and is unique to the large bowel. In contrast, in cases of bacterial overgrowth into the small bowel, normal physiological functions are disrupted resulting in a distinct pathological state. Bacterial overgrowth syndromes of the small bowel, though multifactorial, involve significant interaction of resident bacteria and their byproducts with the gut mucosal epithelial cells. For example, tropical sprue is a syndrome associated with substantial colonization of the proximal small bowel with coliform bacteria that are normally harmless when present in the large bowel (7).

The striking difference of the response to microfloral colonization between cells lining the small bowel and those lining the large bowel indicates the importance of immunological tolerance exhibited in the large bowel. It is a unique functional role of colonic epithelial cells to allow a set of normal microbiota to colonize the lumen of the large bowel, creating a dynamic equilibrium between the degree of microbial colonization and the rate of mucosal renewal.

Conventional wisdom maintains that the rate of renewal of the gastrointestinal mucosa is tightly controlled by the regulation of cell proliferation and programmed cell death by apoptosis. This concept is considered the central dogma of how tissue homeostasis is maintained under normal physiological conditions in the mucosa (8,9). Epithelial regeneration originates from stem cells located in the proliferative compartment comprising the lower third of the colonic crypt. A stem cell may undergo symmetric division to maintain crypt homeostasis, or asymmetric division in which one daughter cell undergoes functional differentiation, migrates toward the lumen and reaches terminal differentiation at the luminal surface (10). These terminally differentiated cells attain “senescence” and are then exfoliated into the fecal stream (9,10). Until recently, little was known about the fate of exfoliated epithelial colonocytes, the generally accepted concept being that exfoliated cells undergo apoptosis or anoikis and were destined to disintegrate during their migration through the alimentary tract.

The investigators of this study however, demonstrated in 1991 that epithelial cells exfoliated into the fecal stream are viable, primarily of colonic origin, and recoverable for further biological studies (11,12). The technology was later refined to enable the isolation of substantial numbers of cells from a small stool sample (0.5 gm) within seven days of collection and following transportation at ambient temperature in a non-toxic preservative solution. We called this procedure “somatic cell sampling and recovery” (SCSR). These cells are exclusively of colonic origin and are pleomorphic in structure and function.

Over a period of eighteen years, SCSR enabled us and other investigators to open a novel approach to the noninvasive study of the pathophysiology of cells originating from the colonic mucosa. Major landmarks were reviewed in a report entitled “Coprocytobiology” to denote the value of this new tool in a distinct field of study opening a new chapter in our understanding of the biological processes in the distal GI tract (13). Exfoliated colonocytes have been studied for biomarkers for mucosal immunity, neoplastic transformation, inflammation and lineage (11-17). SCSR has been used to identify two distinct conditions within the spectrum of inflammatory bowel diseases related to the severity of inflammation with or without a concurrent hyperimmune state (18). We have also demonstrated by RT-PCR that normal, terminally differentiated colon epithelial cells transcribe the genetic information for the insulin receptor, but messenger RNA is not translated into the active protein, in contrast to a reference malignant colon cell line (19).

Using the SCSR technology, other investigators have shown that exfoliated live colonocytes from newborns can serve as surrogates for the evaluation of gut physiology and disease pathogenesis (20). Another study showed that African-Americans (a population known to have an increased prevalence of colorectal cancer) with colonic adenomas expressed a significantly higher proportion of the cancer stem cell markers CD44/CD166 in their exfoliated cell population compared to other groups (21). The transport medium used in this program was recently applied successfully to the room temperature preservation of bovine blood leucocytes *in vitro* retaining viability and functional integrity over a period of eight days (22).

Since the mucosa of the gastrointestinal tract is a major site for the elaboration of immunological defenses mediated by immunoglobulins, SCSR cells were analyzed by flow cytometry for expression of immunoglobulins. These studies revealed a new subpopulation of cells coexpressing antigenic determinants that recognized antibodies to both human IgA and IgG (23). This chimeric immunoglobulin, IgC, appears to be a novel form without a previously described parallel in cell biology. Further studies revealed the existence of immunocoprocytes that expressed only IgA, perhaps representing those mucosal cells involved in the transport of polymeric IgA from the lymphoid cells of the lamina propria. The presence of this complex of cells expressing either IgA or a chimeric IgA/IgG heterodimer presents an interesting insight into the functional specialization within the cells of the mucosa. In this respect, it is also important to note that we were unable to detect significant numbers of cells expressing IgG alone, a distinguishing characteristic of these immunocoprocytes (23).

The present study was initiated to partially map the antigenic profile of cell surface markers on these exfoliated cells. The rationale for this approach was based on the fact that cell surface markers may be masked when they are still anchored on the mucosa and are not amenable to histochemical probes. This was supported by our finding that secretory component (SC), a constituent of secretory polymeric IgA, is undetectable in histochemical sections, yet is readily quantified by flow cytometry on the exfoliated colon epithelial cells. Our survey of the colon cell surface was constrained by the limitations imposed by the available resources and therefore limited to a select number of antigenic markers. The selection criteria was based on a potential application to a disease receptor for which no other approaches are available, (e.g. CXCR-4, a cytokine coreceptor for HIV 1) or markers that are useful in delineating the status of mucosal immunity or markers related to lineage, inflammatory response or presence of gastrointestinal progenitor stem cells (GIP-C). Flow cytometry was our preferred tool in contrast to others such as RT-PCR because the latter did not always reflect the presence of translated functional proteins on the cell surface as revealed by our earlier work on the insulin receptor (19). We were also interested in probing further into the nature of an observed dichotomy in size of these exfoliated cells. Our preliminary experiments showed the presence of two distinct cell populations based on their size distribution (2.5-5 μm versus 5-8 μm). In this report we propose a sentinel mechanism involving the dedifferentiation of exfoliated colon epithelial cells to cystic progenitor stem cells (GIP-C) expressing the intestinal transit stem cell markers Lgr-5 and Musashi-1(Msi1).

## EXPERIMENTAL PROCEDURES

All authors had access to the study data and have reviewed and approved a draft of the final manuscript, with the exception of GK (deceased prior to completion of study).

### Stool collection and transport

Small aliquots of stool were collected by anonymous volunteers, with age, sex and race provided as identifiers. The samples were collected after having obtained informed consent on the basis of an approved IRB at the Beltsville Human Nutrition Research Center of the US Agricultural Research Service. A 0.5–0.8 gm aliquot of the stool sample (either freshly collected or from a stool repository) were placed in a collection tube containing the transport medium provided in the stool collection kit (SCSR-010, NonInvasive Technologies, Elkridge, Maryland) (Fig.1). The collected samples were sent to the laboratory via courier without refrigeration.

**FIGURE 1.**
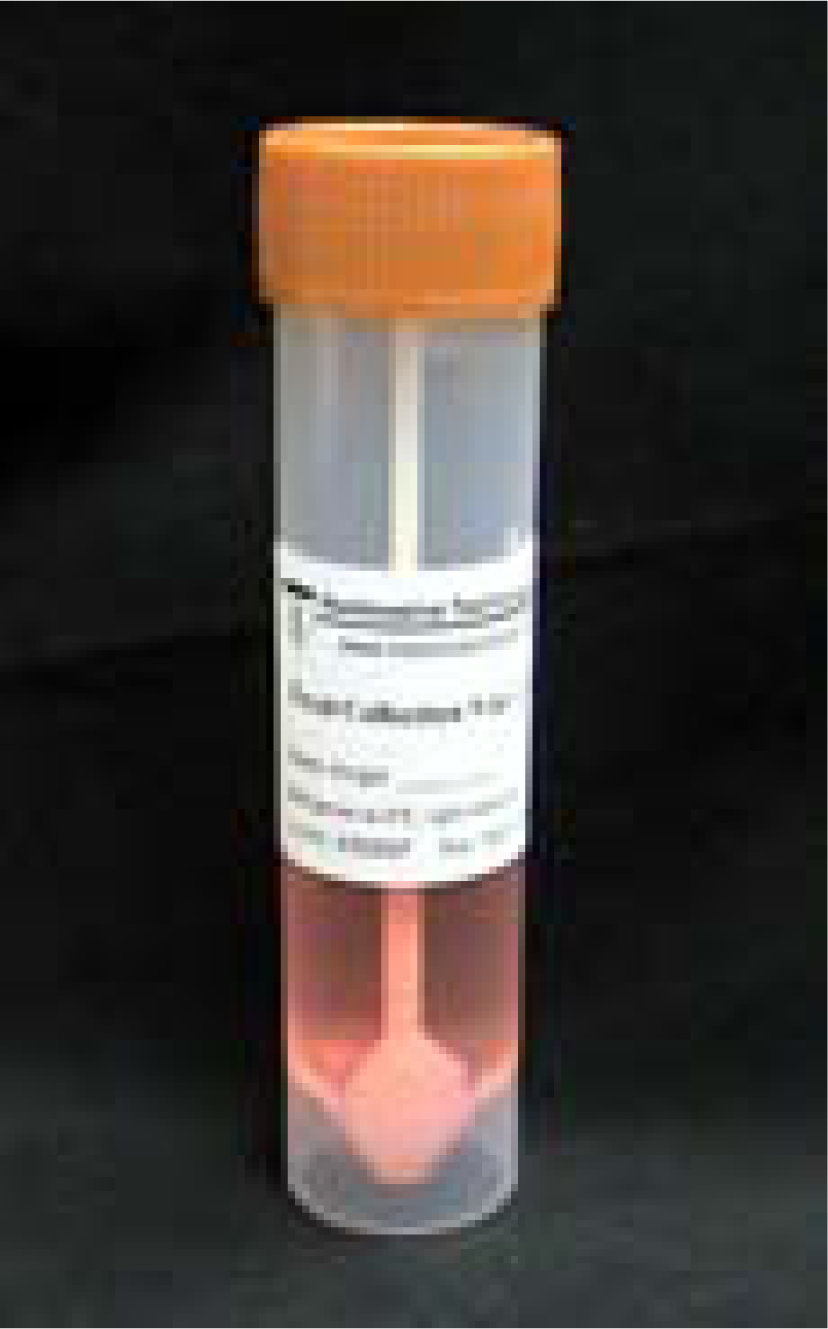
Sample collection tube

### Characteristics of the stool transport medium and field tests establishing performance and stability

The transport medium was formulated to satisfy criteria established during the developmental phase of this program. The composition of the cell isolation medium was designed to allow exfoliated cells to dissociate from the mucus strands to which they were attached while maintaining viability. The medium was stabilized to retain its physiological pH and iso-osmolality. The components of the medium were non-toxic and non-hazardous as defined under US regulations 29 CFR 1910 1200AppA. The system was also designed to protect the exfoliated cells from disintegration while being transported at ambient temperature to the laboratory from a distant site of collection. In addition, the medium arrested replication of the commensal microflora and prevented microbial enzymes from being activated. The final product was filter sterilized and packaged in 10 ml aliquots in sterile collection vials containing about four glass beads (to break up the stool sample) and a secure screw cap to which a long collection spatula is attached (Fig 1). During transport, the collection vials containing about 0.5 – 1.0 gm of fresh stool sample were encased in a larger reinforced plastic cylinder (Sarstedt) to prevent breakage of the sample vials. Following this procedure the viability of cells exceeded 85%. Any deterioration of the transport medium (protected from exposure to ambient light) was easily detected by a change in color of the solution from pink/orange to yellow. Using this criterion we have observed a minimum shelf life of six months.

After having tested the system in the United States, a similar field trial was conducted under tropical conditions. Over a period of two years, approximately 310 samples were collected in Bhuvaneshwar in the Indian state of Odisha and shipped without refrigeration to a laboratory in Trivandrum, Kerala. With the exception of one reported spillage in the laboratory, all other samples yielded cells that met all of our established criteria. Under these conditions the collected samples remained in transit for approximately six to seven days with no evidence of loss of cell viability or structural integrity.

### Isolation of colonocytes from fecal samples (SCSR Protocol)

Laboratory personnel were required to wear protective gowns, gloves and mask prior to entering the room for cell isolation procedures. All initial manipulations were carried out in a certified laminar flow hood. Fluid waste was poured into a closed container holding about 100 ml of 10% bleach, and disposable plasticware were committed to a designated biohazard bag lining a stainless steel receptacle. Laboratory waste from these operations was strictly confined to a special area from which they are removed by contractors certified to handle and dispose biohazard materials. These protocols and practices were reviewed by second level supervisors to ensure adherence to these procedures.

The weight of each sample collection tube was measured and marked on the tube prior to distribution to the subjects for stool collection. Collection tubes returned to the laboratory were re-weighed to determine the weight of the stool sample in the container. This process ensured that the stool sample was within the stipulated weight range of 0.5 – 1.0 gm. Sample sizes above this weight range were above the threshold for successful downstream procedures for isolation of cells.

The samples were diluted with an additional 5–10 ml of the SCSR transport medium (SCSR-T, NonInvasive Technologies, Elkridge, Maryland) and vortexed to disperse the stool to allow the cells to be dissociated from their mucoid matrix. All subsequent steps were carried out as represented in Fig. 2. The suspension was filtered through a 330μm strainer filter bag (Whirl-Pak #B01385WA) followed by a second filtration through a 40μm filter cup (Becton-Dickinson #352340) placed over a 50 ml centrifuge tube. The volume was adjusted to 25 ml with transport medium and underlaid with 10 ml of cushion (SCSR-C, NonInvasive Technologies) prewarmed to ambient temperature. The tubes were balanced and centrifuged in a swinging bucket rotor at 200x*g* at room temperature for 10 minutes.

**FIGURE 2.**
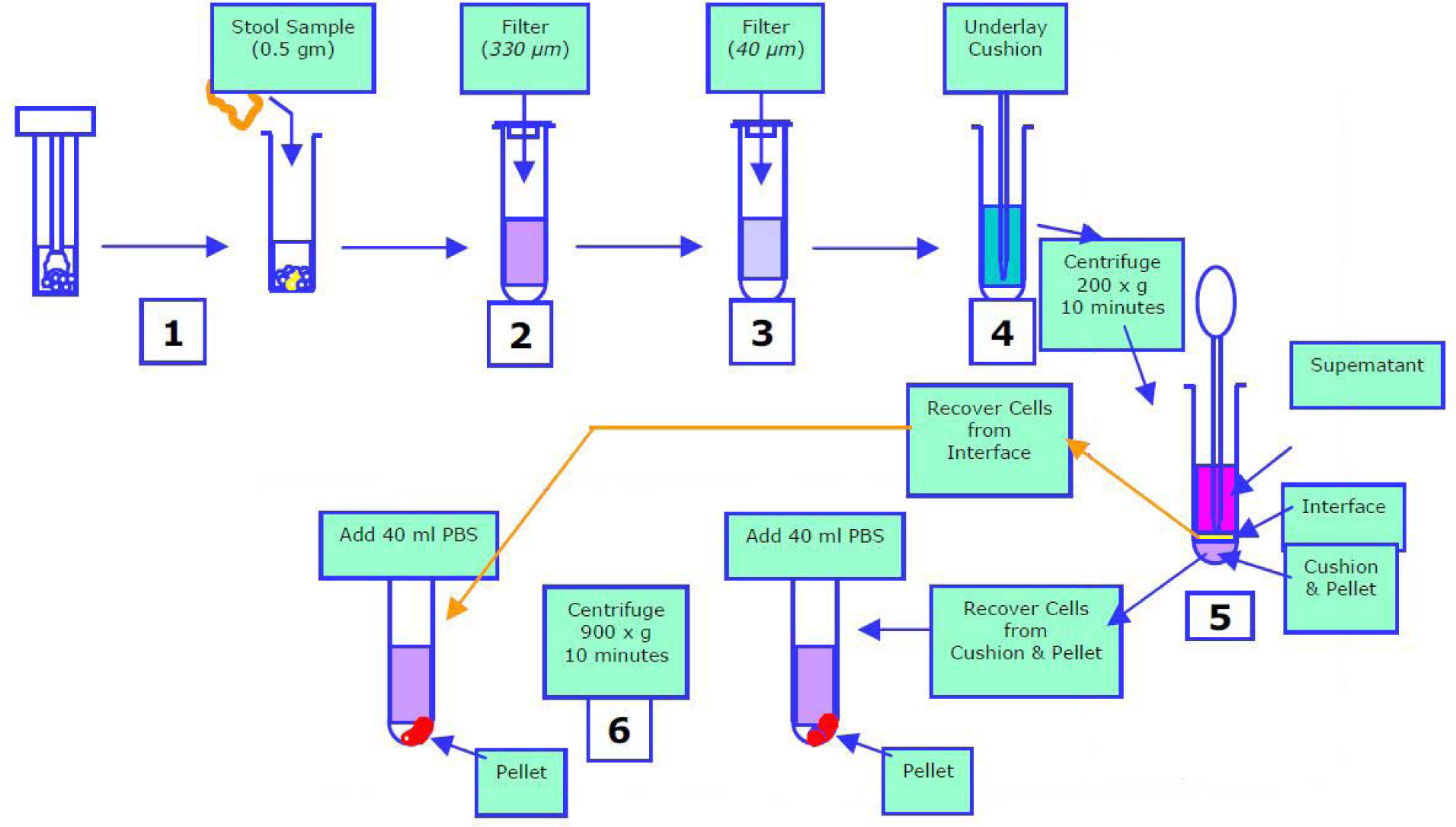
Diagrammatic representation of SCSR cell isolation procedure

The supernatant was removed and discarded into 10% bleach in a waste container. The interphase was carefully removed and diluted to a total volume of 40 ml with cold PBS (pH 7.2) in a second centrifuge tube. The tubes were centrifuged at 900x*g* for 10 minutes at 4°C and the pellets were dispersed in 40 ml of the same buffer. This process was repeated three times. The final pellet was suspended in about 1.0 ml of PBS and counted twice in a Coulter counter, at 2-5μm and 5-8 μm settings. Aliquots of the cell suspension were made according to the desired cell counts and either stored in freezing medium (Sigma #C6295) at -80°C or directly used for experiments on oncolytic activity.

Although newborn infants were not included in the present study, it should be noted that separate studies that did contain infant subjects utilized a modified protocol. In these cases, we initially filtered the stool suspension through a double layer of cheese cloth to remove the interfering meconium and casein. The majority of these cells measured at 2.5 – 5.0 μM, the stipulated range for GIP-C cells. A smaller subpopulation of cells were in the size range of 5.0 – 8.0 μM and were identified as goblet cells by immunocytometry.

### Assay for apoptosis

The percentage of exfoliated colonic cells that enter the apoptosis cycle was determined using annexin V binding in a flow cytometric assay (24). The assay is based on the principle that cells undergoing apoptosis lose their phospholipid symmetry on the plasma membrane (25). As a result phosphatidyl serine translocates to the outer surface of the cell membrane and is readily detected by the specific affinity for annexin V in the presence of calcium ions. This assay was carried out using an apoptosis detection kit (R&D systems, Cat. No. KNX50). Fluorescein-labelled annexin V binding was used to determine the number of cells that bound to the test agent.

### Cell-mediated cytotoxicity against DIO-labelled (Dioctadecyloxacarbocyanine perchlorate, Molecular Probes, Inc) LS-180 adenocarcinoma cells

Dioctadecyloxacarbocyanine perchlorate (DIO), 2.5 mg/ml was prepared in DMSO and 80μl of this solution was added to 20 x 10^6^ LS-180 adenocarcinoma cells suspended in 20 ml of MEM in a T75 Falcon tissue culture flask and incubated overnight at 37° C in a CO2/O2 incubator. The next morning the cells were scraped from the flask, washed in the same culture medium and counted in a hemocytometer. The volume of the suspension was adjusted to give a cell count of about 10^6^ cells/ml.

Serial dilutions of the original SCSR suspension of colonic cells were made in cold PBS to give concentrations of 0.2 – 6.0 x 10^6^ cells/ml. In a typical experiment, a set of six tubes were set up with each tube containing 0.5 x 10^6^ pre-stained LS-180 target cells in 0.5 ml of MEM. Different effector/target ratios were obtained by adding 0.5 ml of the respective dilutions of colon SCSR cells in PBS. Cell suspensions were incubated for 30 minutes at 37°C in a CO2/O2 incubator and then centrifuged at 250x*g* for 5 minutes. The pellets were resuspended in 0.5 ml of PBS. For detecting dead cells, 50μl of propidium iodide (5mg/ml in PBS) was added to each tube and mixed thoroughly.

The suspension in each tube was analyzed in a FACSCAN flow cytometer (Becton Dickinson) optimized for unlabeled LS-180 cells. The scatter gates were set to eliminate any debris making sure that the cells of interest were not excluded from the gates. The log of FL-1 (green fluorescence) versus the log of FL-2 (red fluorescence) was recorded. The percent cytotoxicity was calculated using the following equation:

Number of dead cells/(Number of dead cells + Number of live cells) x 100

### Flow cytometry for surface antigenic markers

Exfoliated colonocytes (both 2.5 – 5.0 micron and 5.0 – 8.0 micron subpopulations) were suspended in PBS 1% BSA at a concentration of one million/ml. Aliquots of 100K were used for the staining and subsequent flow cytometric analysis.

CXCR-4, the chemokine coreceptor for HIV-1, was stained with anti-human CXCR-4 PE (BD Cat. 555974). We chose this marker for its potential value as a tool for the noninvasive detection of latent HIV infection in population surveillance programs. The CD-20 marker for B lymphocytes was stained with anti-human CD-20 FITC (Sigma Cat. C8080). We used this marker to examine the extent of co-expression of CXCR-4 with cells of true lymphoid lineage. The presence of membrane-bound IgA and IgG and a chimeric IgA/IgG heterodimer in a sub-population were identified using anti-human IgA FITC (alpha chain specific, Sigma Cat. F2879) and anti-human IgG PE (Sigma P8047). These markers represented the immunocompetence of the mucosa and test whether there is a consistent age-related senescence in the expression of these markers. The leucocyte common antigen CD-45 was identified using monoclonal anti-CD-45 PE (clone LB-2 PE conjugate, BD Cat 347977). COX-2, as a measure of inflammatory activity in normal subjects was determined using our own FITC-labelled monoclonal antibody raised against an 18-mer polypeptide conjugated with horseshoe crab hemocyanin. Coexpression studies of COX-2/CD-45 were conducted to elicit the ontogeny of cells expessing the inflammatory marker. All flow cytometric analyses included the appropriate isotypes to control for nonspecific binding of antibody to antigen. These values were generally less than 5% and these were subtracted from the test data to obtain the true value for the surface antigen.

### Flow cytometry for Lgr5 and Musashi-1

Since these exfoliated cells readily replicated in culture, they were examined for the expression of Lgr5 and Msi1. Approximately 25,000 colonic SCSR cells were suspended in 1.0 ml of PBS and centrifuged at 900x*g* for 5 minutes. The pellet was suspended in 2.0 ml of 1% BSA in PBS and centrifuged. This step was repeated twice and the final pellet was suspended in 100μl of 1% BSA in PBS. The cells were stained with 5μl of mouse anti-human Musashi-1 (Neuromics, Inc). The cells were incubated at room temperature for 15 minutes, then diluted with 1% BSA in PBS and centrifuged at 900x*g* for 5 minutes at 4^o^C. The supernatant was discarded and the pellet rewashed once. The cells were then stained with a secondary anti-mouse FITC labeled antibody. The stained cells were washed twice with 2.0 ml of 1% BSA in PBS as described earlier and then subjected to flow cytometric analysis along with the appropriate isotype control. This procedure was repeated with cells stained with anti-Lgr5 (GeneTex, Inc). Because of the wide dispersion in size of the SCSR colonocytes, a region gate based on the unstained autofluorescence plot was used instead of a quadrant gate.

### In vitro culture of SCSR exfoliated cells

Two growth media were used for culturing exfoliated cells. In the first system we used Mesencult™ basal medium for human mesenchymal stem cells (STEMCELL TECHNOLOGIES, INC. Cat. No.05401) mixed with mesenchymal stem cell stimulatory supplements (STEMCELL TECHNOLOGIES INC., Cat. No. 05402). In the second system we used Methocult™ SF^BIT^H4436 (STEMCELL Technologies, INC., Cat. No. 04436). The results were comparable in both instances.

About 1-5 million exfoliated cells (2-5 micron Coulter counter window) were used for each batch. Frozen samples were washed 2-3 times with PBS to remove the freezing medium prior to placing them in the growth medium. The cell suspension was passed through an 18 gauge needle 8 - 10 times to breakup mucous strands and dissociate cell clumps to form a single cell suspension. The cell suspensions were initially sedimented in a centrifuge at 2000x g for 10 minutes and the cell pellet was suspended in 2 ml of either growth medium. The cells were aspirated by gently pipetting back and forth a few times before transfer into 6-well plates. The plates were incubated at 37°C in a 5% CO_2_ environment. Cells typically remained in suspension. After approximately one week, the colour of the medium turned yellow. Rather than discarding these samples we waited for the medium to change back to a pink colour (1-2 weeks). After this point the cells were recovered, washed in PBS and subcultured. Optimum results were obtained when the cells were sub-cultured once a week with the subculture ratio set at 1.2 – 1.3. At each passage an aliquot of the cell suspension was dispersed in a serum-free freezing mediuim (Sigma Cat. No C 2639) and preserved in liquid nitrogen.

These cell cultures appear to thrive in the presence of the usual commensal microbiota of the human gut, unlike in other traditional cell culture techniques. In order to prevent contamination of other cell cultures being carried out in the same facility, we grew these cells in a separate CO_2_ incubator in a separate room. Over a period of several years we have consistently reached a passage number close to 90.

Attempts at suppressing bacterial growth with antibiotics repressed the ability of the “stem cells” to replicate, suggesting a crucial role played by commensal microbiota in the maintenance of the replicative potential of these cells.

## RESULTS

### Exfoliated cells are recoverable in a viable state

SCSR technology consistently yielded several million cells per gram of stool, wet weight. Size distribution studies showed the presence of two distinct cell populations, one in the range of 2.5-5 μm and a second one at 5-8 μm. There was a preponderance of the smaller cells, of which about 10% upon storing at 4°C for a few hours showed a shift to the larger size.

### Only about 10% of the exfoliated colonic cells undergo apoptosis

Viability of exfoliated colonic cells isolated from fecal samples (“SCSR Cells”) was assessed by trypan blue exclusion (85%), an observation that was confirmed by flow cytometry using propidium iodide exclusion. In order to assess the status of apoptosis, these cells were examined for binding to annexin V by flow cytometry (24) (Fig 3).

**FIGURE 3.**
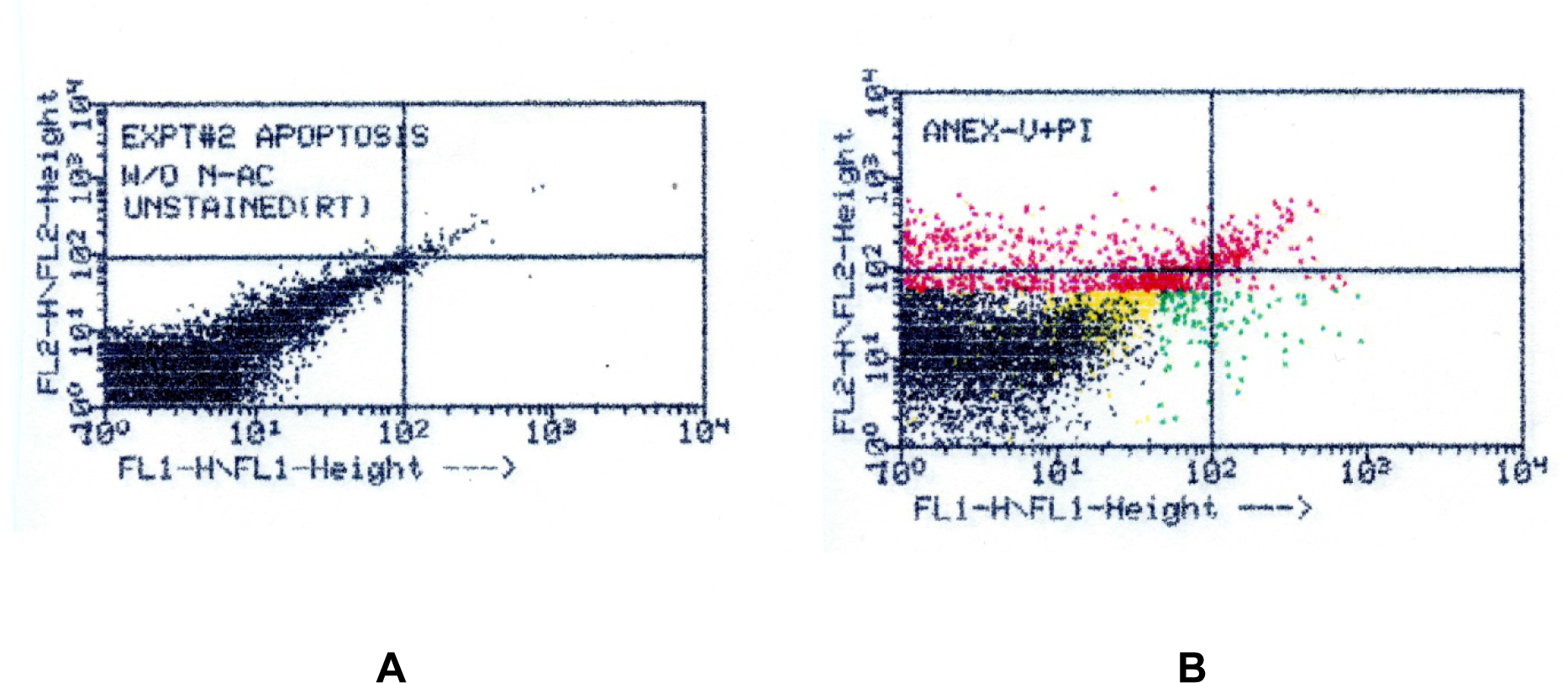
Flow Cytometry – Annexin V A. Isotype control B. Annexin V binding shown in the upper left quadrant

The conventionally accepted concept is that these exfoliated cells are terminally differentiated, partially senescent and undergoing apoptotic cell death (25). Our results seem to be at variance with this assumption since almost all of the SCSR cells were viable by two independent criteria, and no more than 10% exhibited apoptosis as revealed by annexin V binding in FACS assays.

### Cells isolated by the SCSR procedure recognize antibodies directed against colon-specific antigen

We examined exfoliated cells by flow cytometry after treating them with a fluorescently-labeled antibody against colon-specific antigen (CSA) (26). Data from our studies clearly established that exfoliated cells isolated from stool samples are exclusively of colonic origin (13). Further investigations using organ-specific antibodies revealed that a majority of the larger cells (5–8 μm) were primarily goblet cells (David Gold kindly performed these studies using his proprietary organ/cell-specific antibodies) (27).

### Partial map of cell surface antigens

Table 1 presents data on the expression of cell surface antigens from a cohort of 54 subjects. Statistical analyses were performed on a number of parameters from which we have presented the results of gender and race. Age associated expression of these markers did not reveal any consistent trend of senescence. CXCR-4, the cytokine co-receptor for HIV-1 was expressed in over 85% of the cells examined. Almost 50% of these cells also coexpressed lymphocyte biomarker CD20, but were negative for lymphoid lineage marker CD45. It would appear that almost all of the CD20^+^ cells were committed to CXCR-4. While there were no differences between gender, there was a small but significant drop in the values among African-Americans.

**TABLE 1.**
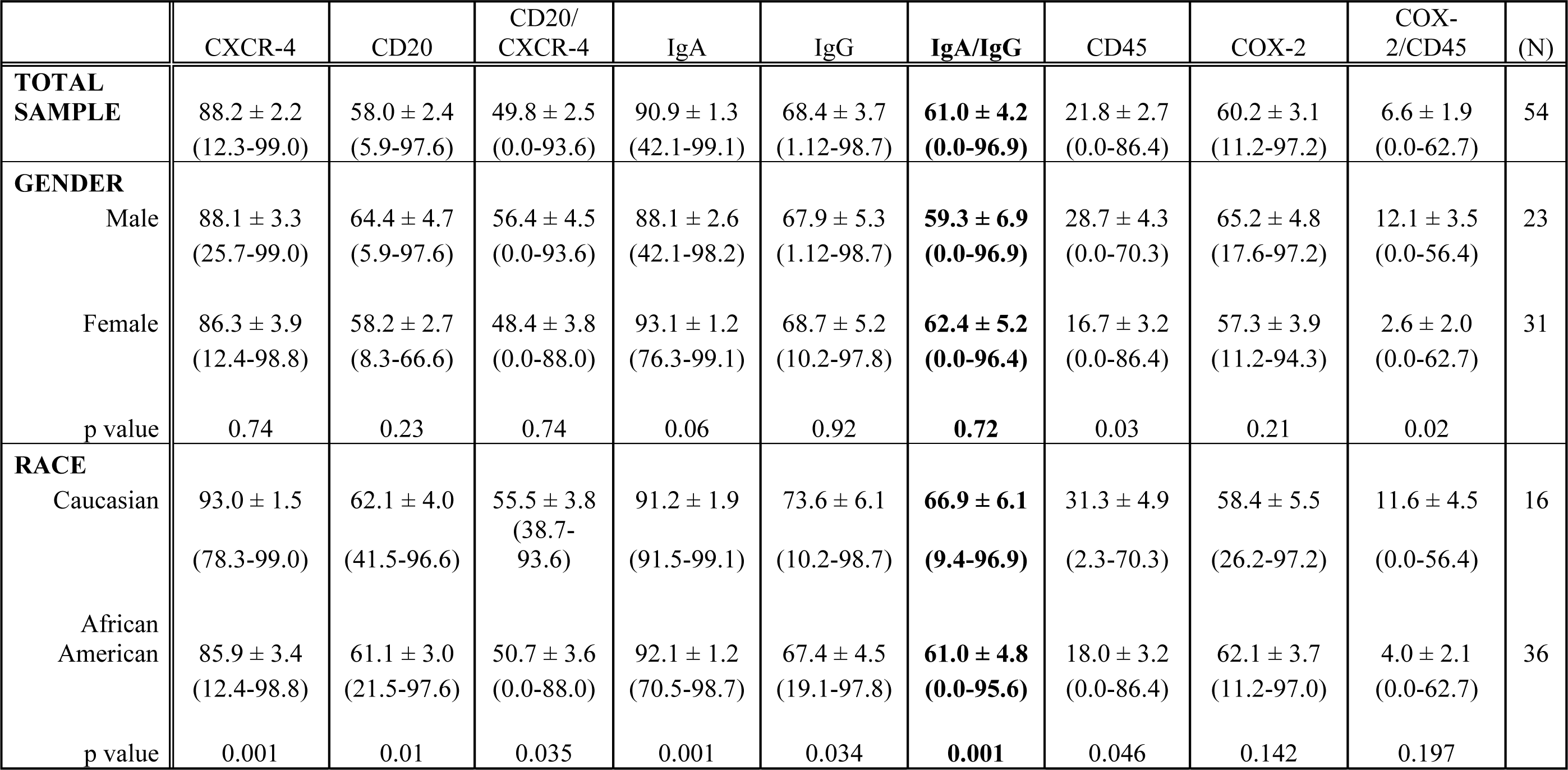
Profile of colonic cell surface markers. Values are means ± SEM; values within parentheses represent the range; CD 20/CXCR-4, IgA/IgG, and COX-2/CD45 are coexpression values by two-color immunofluorescence.

Individually, IgA and IgG were expressed in a majority of cells regardless of gender. Among African-Americans and Caucasians there were significant differences in both IgA and IgG. The most interesting observation was the identification of a subpopulation of cells expressing an IgA/IgG chimeric immunoglobulin that recognized antibodies to both IgA and IgG. Although the presence of this hybrid chimeric antibody (IgC) had been observed earlier, (23) this was the first observation within a large cohort of human subjects. This chimeric immunoglobulin was absent in a small group of African-American subjects. The mean values for the group ranged between 50%–70%. While there were no gender differences, the expression of this immunoglobulin among African-Americans was slightly lower but significant compared to that of Caucasians. The two colour immunofluorescence data for IgC when compared to the single colour data for IgA or IgG alone showed the existence of a significant subpopulation of cells that expressed only IgA while expression of IgG by itself was insignificant. The identification of this chimeric immunoglobulin opens new avenues for investigation on the role of this bispecific antibody. Blocking the IgG component in this chimeric immunoglobulin by prior exposure of the cells to anti-human IgG immunoglobulin discharged the oncolytic activity of these cells showing a potential role of this hybrid in control of aberrant cell proliferation.

CD45, the leucocyte common antigen that signals the presence of cells of lymphoid lineage was seen in a little over 25% of the cells from men, whereas this value dropped significantly in women. Similarly, Caucasians had 30% of the cells expressing this marker compared to only 18% for African-Americans.

COX-2 as an indicator of the standing level of this intermediate in the synthetic pathway for prostanoids revealed a marked presence in these exfoliated cells, indicating an active role for this enzyme although this may also coexist in inflammatory processes (18). The contribution of true inflammatory cells to the expression of inflammatory component COX-2 is further delineated by two-colour immunofluorescence examining its coexpression with CD45. In this cohort of normal subjects we have seen only a small fraction of the COX-2-expressing cells to be of lymphoid lineage (CD45^+^) as would be expected. Samples from women had significantly lower values for inflammatory cells compared to men. This is consistent with our observations that COX-2 is associated with ovulation and that in nonovulatory or postmenopausal women these values tend to be lower or negligible (most of our subjects were postmenopausal). (SL, PPN, unpublished observations)

### Exfoliated colonic epithelial cells exhibit cell-mediated cytotoxicity specifically directed against replicating human tumor cells

After developing the colonic cell isolation procedure we designed a set of studies to determine the cell recovery rate under simulated conditions. In these studies we opted to use the colon adenocarcinoma cell line LS-180, previously labeled with tritiated thymidine as our reference standard.

A second aliquot was processed in parallel as our control reference with only LS-180 cells. The gradient was fractionated and radioactivity-assayed to determine the recovery of the labeled LS-180. In these recovery experiments of labelled tumor cells from a Percoll/BSA linear density gradient as described previously (12) we made the unexpected observation that LS-180 cells in the test sample containing the donor colonic cells consistently sedimented at a density far higher than in the control sample. This “pull down effect” was ultimately shown to be the result of cytolytic activity on the tumor cells leaving what we suspect to be the intact nucleus along with aggregates of adherent colonic cells, accounting for their migration to a denser layer. This indicated to us that exfoliated colonic cells exhibit cell-mediated cytotoxicity, a phenomenon that is generally ascribed to NK cells or cytolytic T-lymphocytes. Our cell isolates from subjects with no evidence of gastrointestinal bleeding were devoid of both CD56^+^ (NK cell) and CD4^+^CD8^+^CD45^+^ T-lymphocytes. We concluded that these exfoliated colonic cells represent a unique population of epithelial cells that, in addition to acquiring progenitor stem cell characteristics, exhibit cytotoxicity directed specifically against tumor cells. Exfoliated colonic cells isolated in our laboratory were given to an outside research center for confirmation of our observations. This group confirmed the cytolytic effect on a number of cancer cell lines using their flow cytometric assay procedure (Salata, Walter Reed Army Medical Research Center). In our laboratory, the exfoliated cells isolated from stool samples selectively killed tumor cells without exhibiting cytotoxicity against replicating non-tumor cells in a flow cytometry procedure. We observed over 80% killing of the target cells in all instances as shown in Fig 4.

**FIGURE 4.**
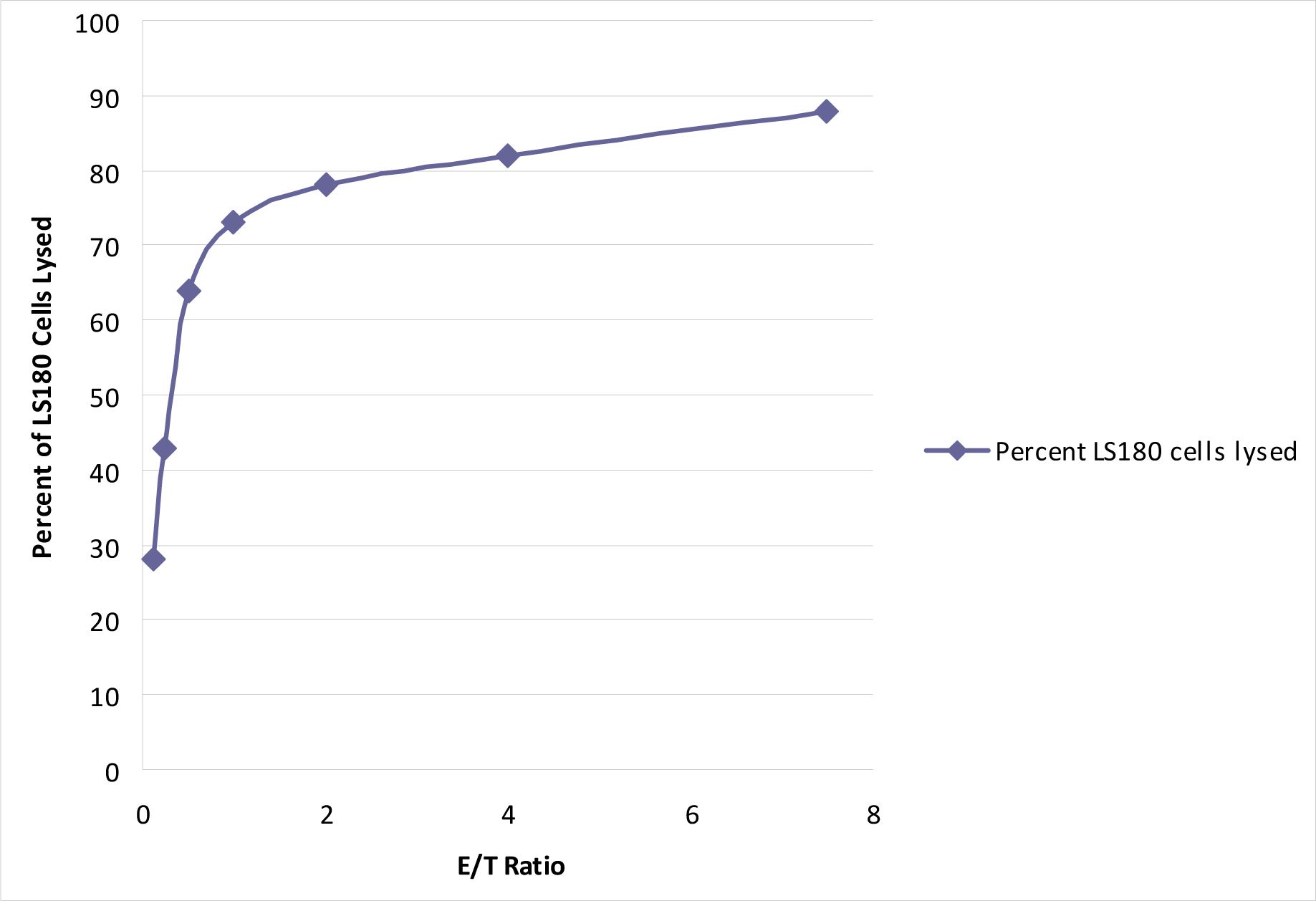
Cell-mediated cytotoxicity of SCSR cells. Plot of effector/target ratio (x-axis) versus percent cell lysis of LS-180 adenocarcinoma cells (y-axis).

The specificity of this activity implied the ability of exfoliated colonic cells to distinguish malignant cells from normal replicating cells. Since pretreatment of these cells to anti-human IgG antibody blocked the cytotoxic effect we are ascribing this effect to the IgG component of the chimeric IgA/IgG molecule in view of the absence of cells expressing only IgG. Cytotoxic T-cells exhibit such specificity mediated via epitope-targeted lineage-specific antitumor activity directed towards the cancer mucosa antigen, guanyl cyclase C (28). However, the inability to detect T-cell markers on our cell isolates does not exclude the possibility of having T-cells in our preparations, as it is known that T-cells can exist as CD4^-^CD8^-^ double negative gamma delta T cell phenotypes exhibiting potent anti-tumor activity (29-32). In this regard there is some evidence that our SCSR cells have a subset of gamma delta T cells. (Weitkamp, personal communication).

### A majority of cells undergo cystic transformation following exfoliation

As phase contrast images suggested that exfoliated cells appear to be smaller than the accepted size of eukaryotic somatic cells (Fig. 5), we subjected them to size distribution analysis (Coulter Analyzer) revealing two distinct cell populations: one in the range of 2.5-5 μm and a second one in the range of approximately 5-8 μm (Fig. 6). The smaller cells appeared to be a distinct population of quiescent cells (several million cells/gram of fresh stool) of which about 10% spontaneously reverted to the larger size following incubation for about six hours at 4°C. This phenomenon of autotrophism, a functionally active process, appears to be a response to an as yet unidentified secretory effector in the milieu. The larger cells were morphologically and immunocytometrically consistent with goblet cells.

**FIGURE 5.**
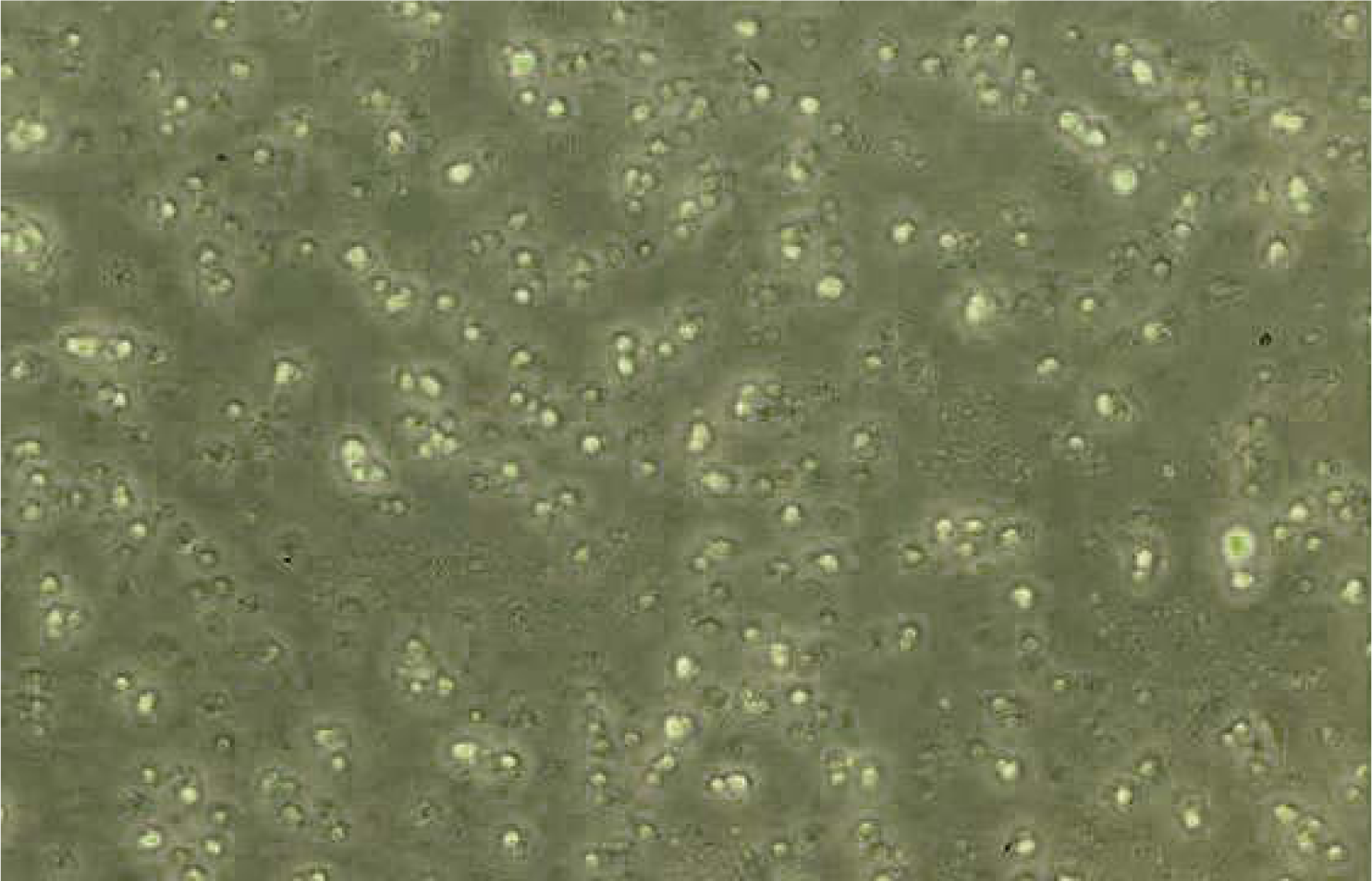
SCSR cells isolated from stool. Phase contrast image

**FIGURE 6.**
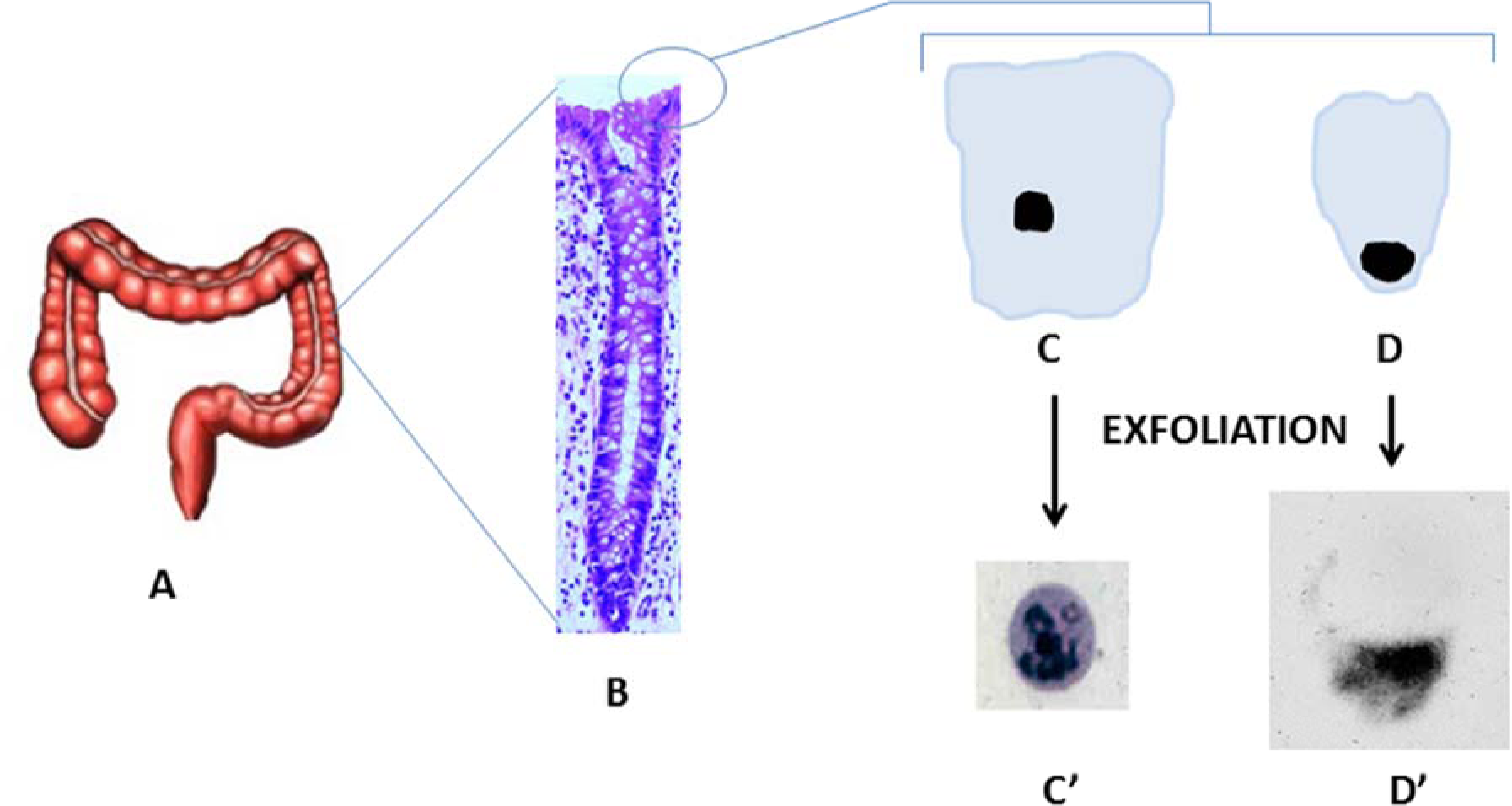
Diagrammatic representation of exfoliation and dedifferentiation of colonic epithelial cells to give rise to cystic GIP-C cells. A. Colon. B. Crypt. C. Columnar Cell. C’. Exfoliated Columnar Epithelial Cell – dedifferentiates into cystic GIP-C Cell expressing Lgr-5 and Musashi-1, approx. 3-4 microns in size (Wright-Giemsa stain). D. Goblet Cell. D’. Exfoliated Goblet Cell – shape and size appears unaltered on exfoliation (goblet-specific immunostain, courtesy David Gold, CMMI, Belleville, NJ)

### SCSR cells replicate in vitro over several generations and can be differentiated into plasma cell-like progeny under controlled conditions

There are numerous reports in the literature describing the in vitro culturing of surface colonic epithelial cells isolated directly from the mucosa of resected specimens from humans and experimental animals (33-36). As early as 1983, Moyer alluded to the presence of stem cells in the central region of replicating cell colonies growing in culture (33). Stool-derived cells can also be cultured, and we have cultured them for over 90 generations. In addition we were able to differentiate them in a lineage directed fashion into plasma cell-like progeny that generate human tumor-specific antibodies. IgG isolated from these culture supernatants arrest tumor growth in xenotransplants (HT-29 human adenocarcinoma) in SCID mice (unpublished observations).These replicating stool derived cells, presumably originating from the “terminally differentiated” epithelial cells, exhibit functional characteristics that fit the quintessential definition of a progenitor stem cell – the capability for long term replication and the amenability to lineage-directed differentiation.

### Exfoliation from the mucosal surface may be accompanied by spontaneous dedifferentiation

It is significant to point out that there are several reports of colonic cells being easily maintained in culture over several generations without regard to the fact that the vast majority of the initial founding cells are nonreplicating, and terminally differentiated according to currently accepted principles (33-36). Therefore, this apparent ability to replicate may be ascribed to a newly acquired characteristic involving dedifferentiation associated with a reprogramming phenomenon, one that has been reported on a restricted basis in other organ systems as part of the normal biology of the living organism (37-42). We believe we are observing a similar reversal of the generally accepted dogma of growth and differentiation in the forward direction in GIP-C cells. Dedifferentiation as a normal biological phenomenon in the rejuvenation of injured or diseased tissue has been championed by Blau and other investigators, and is now being recognized as part of the body’s reconstructive processes (42).

### Dedifferentiation is associated with a transition to a spore-like state

The presence of spore-like stem cells (less than 5 μm) in several tissues have been described by Vacanti and associates (43). Being dormant, they can survive in extremely low oxygen environments and hostile conditions such as extreme temperatures. These are small undifferentiated cells that are predominantly nucleus with a small amount of cytoplasm. *In vitro* these structures have the capacity to enlarge, develop and differentiate into cell types expressing characteristics appropriate to the tissue environment from which they were initially isolated. Vacanti *et. al* (43) believe that these unique cells lie dormant until activated by injury or disease and that they have the potential to regenerate tissues lost to disease or damage.

Shmilovici (44) has reported similar spore-like cells that are small and dormant with simple structures and have the ability to differentiate into mature cells. They are present in every tissue in the body and tolerate conditions that kill differentiated or partially differentiated cells, such as complete oxygen deprivation and exposure to temperatures much higher or lower than normal body temperature. When activated, they can proliferate more rapidly and into more types of differentiated cells than terminally differentiated or stem cells. Shmilovici’s hypothesis is that they provide redundancy for the aging cells in the tissue.

Recent studies have shown that the “terminally differentiated” state of human somatic cells is not irreversible, and can be altered while concurrently inducing the expression of previously silent genes associated with undifferentiated states (37-42). Even a small molecule, a 2,6-disubstituted purine named reversine, has been shown to induce reversal of differentiated C2C12 cells to become multipotent progenitor cells that can redifferentiate into osteoblasts and adipocytes (41).

### GIP-C cells are spore-like and express the gastrointestinal stem cell markers Lgr5 and Musashi-1

Our work on exfoliated colonic epithelial cells has shown that upon exfoliation a majority of the cells shrink in size (2.5–5 microns) with the plasma membrane condensing along with the cytoplasm over the nucleus to form spore-like cystic bodies (Fig. 6). In spite of this transformation, these cells are functionally active, expressing the putative intestinal progenitor stem cell markers Lgr5 (45) (28%-97%) and Musashi-1(46) (6%-75%), are capable of replication and show plasticity usually associated with stem cells (Figs. 7, 8). Our yield estimates of these stem cells from stool vary, and do not appear to be influenced by age, sex or race.

**FIGURE 7.**
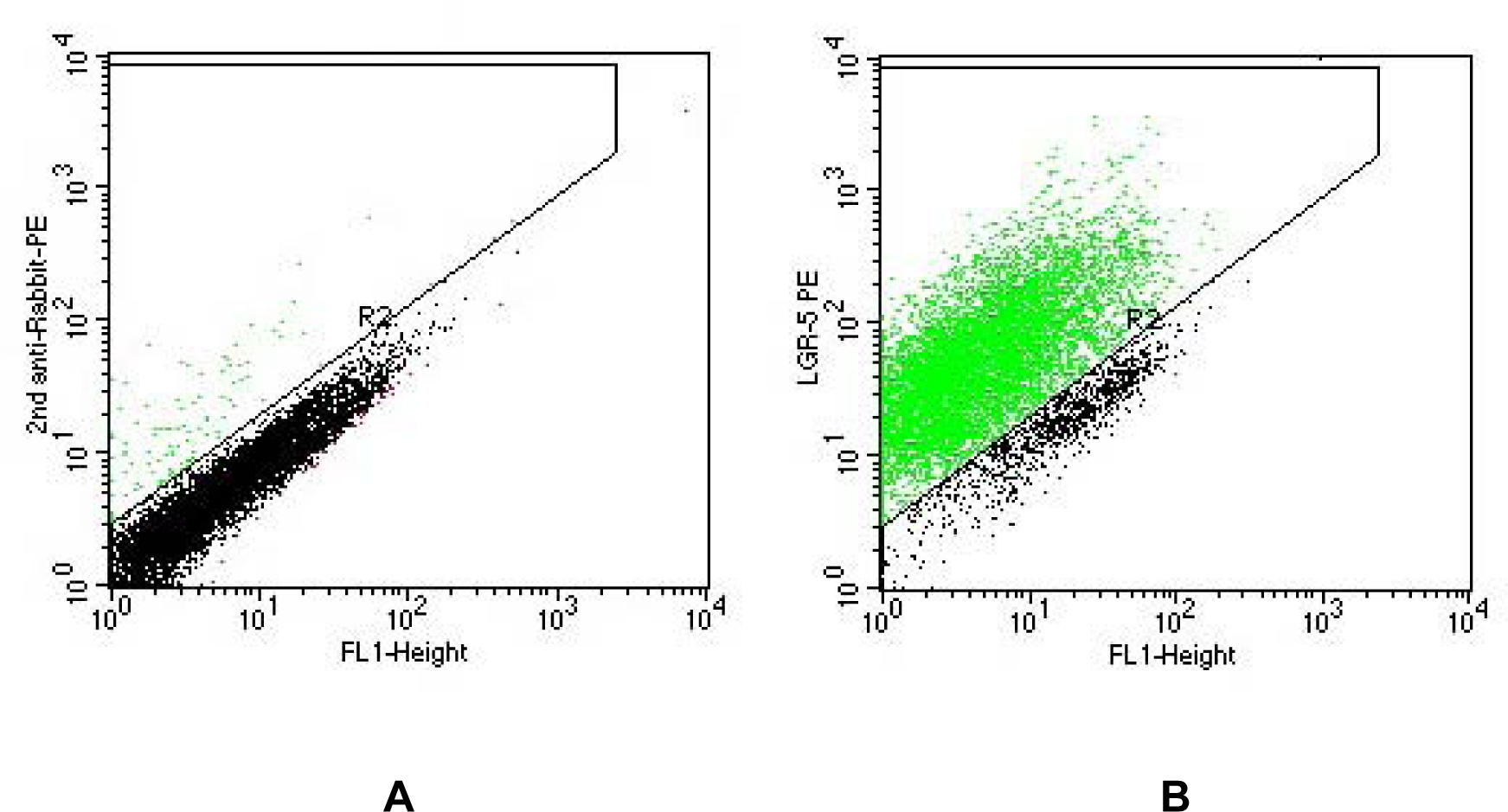
Flow Cytometry – Lgr5. A. Isotype control. B. Lgr5 expression in R2

**FIGURE 8.**
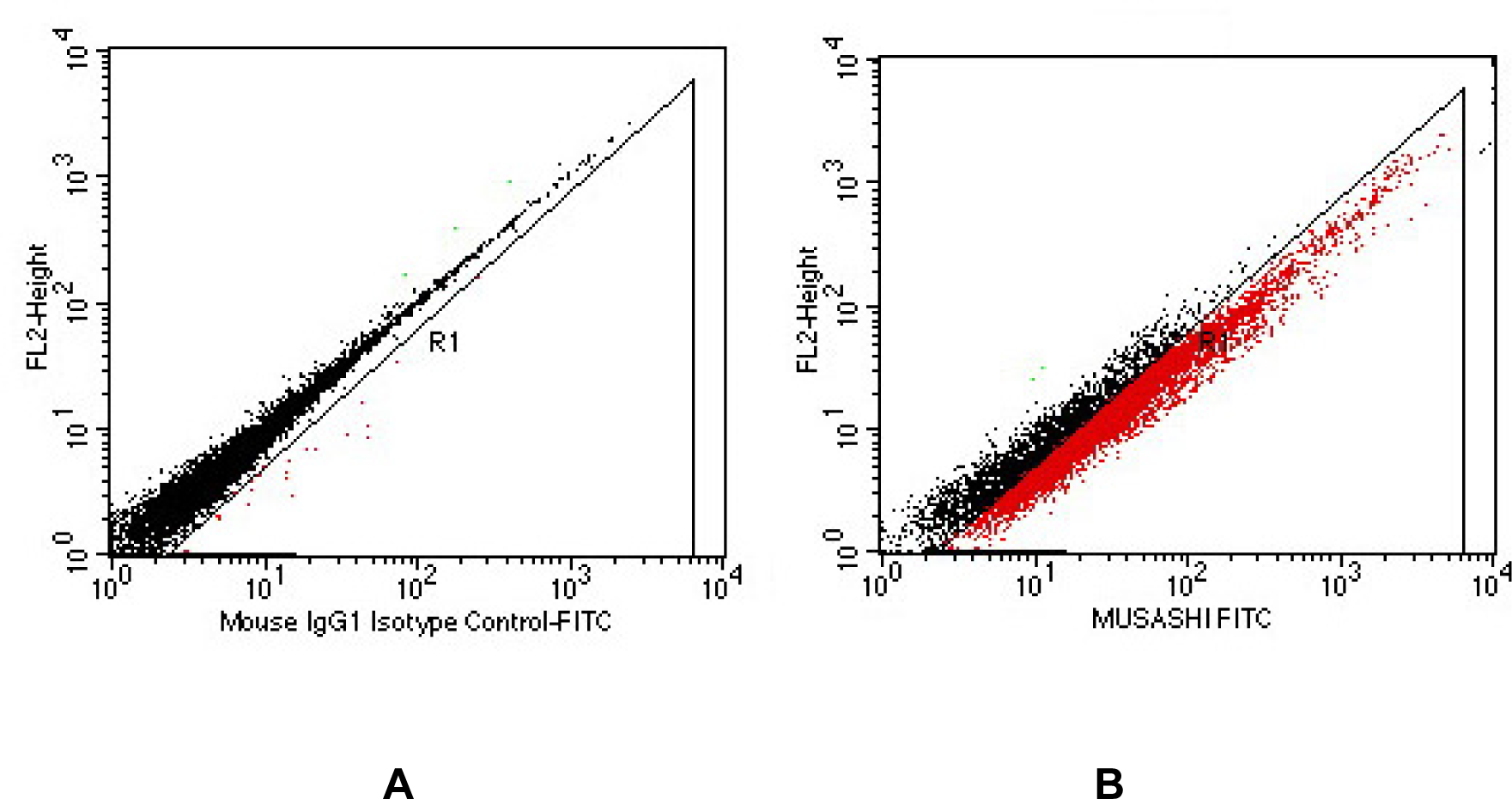
Flow Cytomery – Musashi-1. A. Isotype control. B. Musashi-1 expression in R1

## DISCUSSION

There is a significant body of evidence relating to the expression of cytoskeleton and differentiation markers in the colon. Some of the early work in this area was reviewed in a comprehensive report by Ho (47). Almost all of this knowledge was derived from histochemical and immunofluorescence studies, where the observations were limited by the masking of the markers or restricted access to the cellular surface by the tightly packed structure of the colonic epithelium. During the course of two decades of investigations into the nature of cell surface markers we found evidence to support the concept that exfoliated colonic cells may be undergoing remodeling as revealed by the expression of Lgr5 and Msi1. The appearance of a cohort of cells associated with the coexpession of IgA/IgG is attributed to their denovo synthesis as a heterodimer and not as a nonspecific binding of the two entities to cell surfaces. There are two lines of evidence to support this assumption: One is that the cell collection medium is formulated to dissociate any molecules that adhere nonspecifically to cell surfaces. The second is that the cells when cultured consistently generated the IgA/IgG heterodimer in the progeny showing the endogenous origin of this immunoglobulin. The total absence of this subpopulation exclusively in a small number of African-Americans point to a germline deletion of this signal. Hsu (48) has described the *de novo* coexpression of multiple immunoglobulin isotypes on human lymphocytes in specific clinical conditions but not in a multimeric form. Our evidence indicates that a study of markers on exfoliated colon cells may be a useful approach to investigate the immunologic consequences of interactions among dietary components, the cells from the epithelium and the commensal microbiota.

## RATIONALE FOR EXPERIMENTAL APPROACH

We selected flow cytometry as our primary tool as opposed to RT-PCR as an example for the detection of functional markers on cell surfaces. This choice was based on our earlier observation that in normal colonocytes the mRNA for the insulin receptor was not translated, resulting in a lack of expression of the functional protein within the cells from healthy subjects (19).

### Cells collected via SCSR represent the anatomic entirety of the colon

In spite of the long transit time for exfoliated cells from the proximal segments of the colon to travel to the anus, we were able to demonstrate that they were recoverable in a viable state (12,13). This was based on the fact that blood group antigens in the adult are localized to cells from the proximal segments (cecum and right colon) of the colon and that SCSR cells expressed this marker proportionately (13).

Our current thinking on the mechanistic aspects for this spontaneous dedifferentiation of large numbers of terminally differentiated epithelial cells rests with the fact that these cells undergo this transformation following exfoliation from a restricted architecture of the mucosa (49). The notion that exfoliation releases mature differentiated cells from the constraints of the structured epithelium is supported by studies showing the dedifferentiation of exfoliated primary human articular chondrocytes (50).

Crypt-based columnar stem cells are replicating cells marked by Lgr5 (45) and Msi1 (46). Their progeny are rapidly dividing transit-amplifying cells that migrate along the crypt axis while undergoing progressive differentiation into specialized structural and secretory cells. By virtue of their location in the deep cryptal segment it is unlikely that a large population of these stem cells are exfoliated into the fecal stream. However, we have shown that up to 90% of the exfoliated GIP-C cells express these gastrointestinal stem cell markers and are capable of being maintained in culture for over 90 generations. Furthermore, as described earlier, they are capable of being converted in a lineage directed manner to plasma cell-like forms generating antibodies against human tumor cell lines, further reinforcing the concept that exfoliation from the colonic mucosa results in a concomitant reprogramming of the terminally differentiated epithelial cells into replicating gastrointestinal progenitor stem cells (GIP-C). (Figs. 7,8)

### Clonal expansion of pluripotent GIP-C cells as a source for therapeutic engraftment

Recently, Sato and Clevers (51) have reviewed work on single intestinal stem cells growing into self-organizing mini-guts, demonstrating the plasticity of these stem cells to reorient themselves into structures mimicking the intestinal crypt. They have emphasized the importance of this observation for potential applications in restitution of the mucosa in gastrointestinal disorders. If this interpretation holds, then we have a clear path to mucosal reconstitution using GIP-C cells in colonic engraftment.

This is further supported by the reported successful treatment of aganglionic gut disorders in humans with engraftment of nervous system stem cells derived from the human gut (52). In such cases, enteric nervous system stem cells were isolated from mucosal colonic tissue obtained by endoscopy and expanded *ex vivo* to generate neurosphere-like bodies. Transplantation of these bodies into aganglionic segments of the human gut resulted in reconstitution of the mucosa, differentiating into enteric nervous system neuronal and glial cells. This demonstrates the plasticity of these colonic mucosal stem cells to engraft onto damaged mucosa and enable restitution of physiological functions following reconstitution into appropriate functional cellular elements of the crypt (52-54).

Intestinal stem cells have been shown to have the innate ability to interconvert between stem cell populations in specific niches within the crypt, demonstrating their plasticity to model the crypt according to the demands of the dysfunctional mucosa. There is also indirect evidence suggestive of bi-directional movement of stem cells along the crypt axis, indicating the ability of these proliferating cells to migrate away from the luminal surface towards the inner proliferating zone where the quiescent and active stem cells are normally anchored.

## CONCLUSIONS

We believe our findings have broad relevance to human disease, as demonstrated by the use of fecal transplant as a new modality for the treatment of intractable *Clostridium difficile* infection (55). This approach has received considerable acceptance in view of the reported efficacy of greater than 90% (56). The beneficial effects have been attributed to the role of the commensal microbiota from the donor stool. Our work shows that fecal transplants include both microbiota and companion viable GIP-C cells, which raises the possibility that the underlying repair mechanism may involve the regenerative engraftment of a new epithelial “lawn”. From the large number of successful fecal transplants without host vs. graft reaction to the allogeneic cells, we believe that these GIP-C cells are of low immunogenicity and readily reseed the host mucosa with epithelial cells that are already accommodative of the transplanted microbiota (57). This interpretation is further reinforced by the fact that orally administered probiotics devoid of epithelial cellular elements do not result in sustained colonization of the gut with beneficial microbiota in cases of *C. difficile* infection (55).

Further support for the concept of regenerative repair of the colonic mucosa by cellular engraftment has been demonstrated by the successful treatment of aganglionic gut disorders in humans (e.g. Hirshsprung’s disease) as stated earlier (52). Therefore, there is compelling evidence to suggest that reconstitution of the colonic mucosa by GIP-C cells present in fecal transplants is a concomitant feature along with the colonization of the associated microbiota that originated from the donor.

In a similar manner, stool transplants from lean donors to obese subjects with metabolic syndrome resulted in improved insulin sensitivity (58). This raises the possibility for the treatment of type 2 diabetes and obesity, which are metabolic diseases associated with insulin resistance and a defect in the production of GLP-2 (glucagon-like peptide 2, part of the incretin effect) by mucosal L cells (59).

Finally, upon reviewing our findings along with the recent observations of the beneficial effects of fecal transplants, we feel that exfoliated colonocytes may play an ancilliary role in defining the pathophysiology of some diseases. The central tenet of our findings is that mature terminally differentiated cells revert to a quiescent cystic dedifferentiated (QCD) stem cell-like state upon exfoliation from glandular epithelium, which can be reactivated extra-corporeally to replicating stem cells and engrafted with or without lineage-directed differentiation. The applications of this approach may be beneficial in the treatment of inflammatory bowel disease (IBD), irritable bowel syndrome (IBS), neurodevelopmental and neurodegenerative disorders, metabolic syndrome, obesity, diabetes, spinal cord injury, and cancer (we have successfully used GIP-C cells for generating tumour-specific antibodies that arrested tumour growth in xenotransplants in SCID mice). We believe that GIP-C cells are a valuable addition to other potential sources of quiescent progenitor stem cells, including neural crest stem cells from the GI tract.

## ACKNOWLEDGEMENTS

This paper is dedicated to the memory of George Kessie who unflinchingly participated in the early stages of this work and conducted some of the exploratory studies that paved the way for the current state of our knowledge We thank Roy K.H. Wong (Walter Reed Army Medical Center) for his keen and long standing interest and advice on the gastroenterological aspects of our work. Pinaki Panigrahi (University of Nebraska Medical Center) facilitated the field trials in India. Steven Czinn (Chairman, Department of Pediatrics, University of Maryland School of Medicine) and Jay Nair (Vanderbilt University School of Medicine) and Prabha Chandrasekharan, (National Institutes of Health) reviewed the manuscript and gave valuable suggestions.

## AUTHOR CONTRIBUTIONS

PPN conceived the program, obtained the funding, directed the study and wrote the paper. GPA, VI, SS, AL, ES, GK, SL, AK, JJC, SK, and RS contributed to the performance of experiments and acquisition of data. PN unblinded data sets and conducted statistical analyses using SPSS and verified performance reliability of the cell isolation technology. GPA, VI, SS, SD, AL, PN, RN and JC assisted in critical review of the manuscript, LEP reviewed and edited the final manuscript.

## Footnotes

** In this article, “SCSR cells” denote exfoliated cells isolated from stool samples and “GIP-C” are progenitor cells either present in SCSR cell isolates or cultured from SCSR cell isolates.

